# Modeling Marek’s disease virus transmission: a framework for evaluating the impact of farming practices and evolution on disease

**DOI:** 10.1101/185785

**Authors:** David A. Kennedy, Patricia A. Dunn, Andrew F. Read

## Abstract

Marek’s disease virus (MDV) is a pathogen of chickens whose control has twice been undermined by pathogen evolution. Disease ecology is believed to be the main driver of this evolution, yet mathematical models of MDV disease ecology have never been confronted with data to test their reliability. Here, we develop a suite of MDV models that differ in the ecological mechanisms they include. We fit these models with maximum likelihood in ‘pomp’ to data on MDV concentration in dust collected from two commercial broiler farms. Using AIC to compare the models, we find that virus dynamics are influenced by between-flock variation in host susceptibility to virus, shedding rate from infected birds, and cleanout efficiency. We also find evidence that virus is reintroduced to farms approximately once per month, but we do not find evidence that virus sanitization rates vary between flocks. Of the models that survive model selection, we find agreement between parameter estimates and previous experimental data, as well as agreement between field data and the predictions of these models, suggesting robustness of model predictions. Using the set of surviving models, we explore how changes to farming practices are predicted to influence MDV costs, should pathogen evolution undermine existing control measures. By quantitatively capturing the mechanisms of disease ecology, we have laid the groundwork to explore the future trajectory of virus evolution.

## Introduction

Marek’s disease virus (MDV), the causative agent of Marek’s disease (MD), imposes a substantial economic burden on chicken meat and egg production, costing the worldwide poultry industry in excess of 1 billion USD per year (Morrow and Fehler 2004). Historical control measures have at least twice been undermined as a result of virus evolution, leading to speculation that future evolution could undermine current control (Nair 2005). The ecology of the disease appears to be the driving force behind past evolution, with explanations invoking vaccination (Witter 1997; Atkins et al. 2013a; Read et al. 2015), rearing period duration (Atkins et al. 2013a; Rozins and Day 2017), and virus persistence during downtime between bird flocks (Rozins and Day 2017). Understanding the ecology of the virus is thus a key component in predicting whether and when control efforts will lose efficacy. Such an understanding is also crucial in developing immediate responses should the efficacy of current control measures wane. Yet the ecology of MDV is poorly understood. This is perhaps most clearly exemplified by the conventional wisdom saying that the virus is ubiquitously found on industrialized poultry farms (Office International des-Epizooties 2010; Dunn 2013), despite recent surveillance data suggesting that the virus may not be present on a large fraction of farms (Groves et al. 2008; Wajid et al. 2013; Walkden-Brown et al. 2013; Bettridge et al. 2014; Kennedy et al. 2015b; Ralapanawe et al. 2015; Kennedy et al. 2017).

Mathematical models of disease ecology can provide valuable insight into infectious disease dynamics. Such models quantitatively relate changes in ecology to changes in disease dynamics, which is particularly useful when experimental manipulation is unethical or, as with commercial-scale chicken rearing, financially costly. Models thus provide cheap and safe opportunities to explore the impact of system manipulation on pathogen control, and this approach has justly been applied to MDV (Atkins et al. 2013a,b; Rozins and Day 2016, 2017). The reliability of a model, however, can only be assessed by challenging it with data, and this has never been done for MDV. Here we develop a suite of models to describe MDV dynamics on commercial broiler farms, and we use model selection methods to identify the ecological mechanisms that are most important to explaining MDV dynamics in the field.

Poultry intended for consumption are inspected and condemned for a condition called “leukosis” at the time of processing. This condition can be caused by various diseases, but in chickens reared for meat, leukosis is almost exclusively caused by MD (Sharma 1985). Current rates of condemnation due to leukosis are extremely low (Kennedy et al. 2015b), but future reductions in vaccine efficacy due to virus evolution might cause leukosis rates to increase, as was documented with the erosion of vaccine efficacy in the past (Witter 1996). A method to relate changes in farming practices to changes in risk of condemnation would therefore be a useful tool should virus evolution continue along the trajectory of the past.

Surveillance data have shown that the concentration of MDV in dust can vary several orders of magnitude between farms and within farms over time (Walkden-Brown et al. 2013; Kennedy et al. 2017). The underlying cause of this variation is unknown. Mechanistic explanations might include between flock variability in virus susceptibility and in virus shedding. Such differences can arise because of choice of bird breed, quality of chicks, and choice and efficacy of vaccines. Mechanistic explanations might also involve differences in husbandry, such as differences in the efficiency of virus removal, in the sanitization efficacy in houses, or from reintroduction of virus (biosecurity). By comparing mathematical models that include or exclude these various mechanisms, we can identify the importance of these differences on MDV dynamics, and in turn, we can develop strategies to control the ecology, evolution, and economic burden of this pathogen.

## Methods

### Model construction

We model the transmission and persistence of MDV within and between flocks of broiler chickens on commercial poultry farms. Our models are constructed assuming standard rearing practices in Pennsylvania, United States, practices which are fairly standard for commercial poultry rearing across the United States and much of the developed world.

Industrial-scale rearing of broiler chickens on farms tend to follow an “all-in, all-out policy,” meaning that all chickens within a house are reared as a single-aged cohort of birds. We refer to this cohort of birds that occupy a single house on a farm as a flock. Birds are placed on litter that consists of wood chips and sawdust at one-day-old, and the birds are reared in this environment until they are ready for processing. Birds in houses are provided ab libitum food and water. Temperature, humidity, and air quality are maintained by a combination of active ventilation through fans or wind tunnels and heating. Flocks are collected for processing when sufficient time has elapsed for birds to reach a particular target weight.

While chickens are being reared, houses accumulate “chicken dust,” a by-product of farming that consists of bits of food, epithelial cells, dander, bacteria, and feces (Collins and Algers 1986). The amount of dust produced by birds increases as birds grow (Islam and Walkden-Brown 2007; Atkins et al. 2013a). Infectious MDV can be contained in this dust (Carrozza et al. 1973), being shed with the epithelial cells of infected chickens and transmitted through the inhalation of virus-contaminated dust (Colwell and Schmittle 1968). The concentration of MDV in dust can be measured through quantitative polymerase chain reaction (qPCR) (Baigent et al. 2005; Islam et al. 2006; Baigent et al. 2016). Our model is constructed with this type of data in mind. We thus track the infection status of birds as well as total dust and total virus quantities.

Coming from extremely hygienic hatcheries, chickens are unexposed to MDV when first placed in a house. Shedding of virus from a bird can begin as early as one week post exposure to virus and tends to reach maximal levels two to three weeks post exposure (Islam and Walkden-Brown 2007; Read et al. 2015). Once reached, virus shedding stabilizes at peak levels for the duration of a broiler chicken’s life (Islam and Walkden-Brown 2007; Read et al. 2015). Shed virus can infect other chickens, causing the pathogen to spread to other hosts in the flock.

Typical commercial broiler farms vaccinate against MD by using bivalent vaccination (Morrow and Fehler 2004). Although vaccinated birds can still be infected with MDV and can still shed MDV (Witter et al. 1971; Islam et al. 2008; Ralapanawe et al. 2016), vaccination greatly reduces clinical signs of disease (Witter et al. 1971). This, along with other measures to ensure bird health, means that total morality from hatch to processing is typically minimal (≈ 3% and ≈ 8% in the two farms used for model inference later, unpublished data – in line with the national average of 4.8% (National Chicken Counsil 2016)). We therefore assume that bird mortality is negligible in our model. A schematic representation of the infection dynamics are shown in fig. 1, corresponding to the following set of mathematical equations:

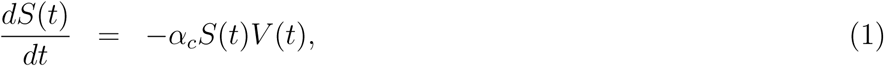

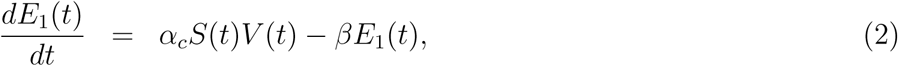

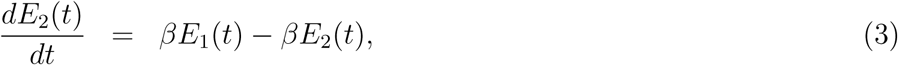

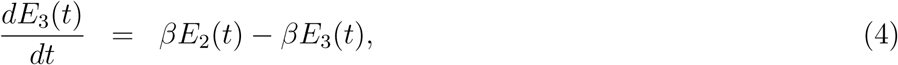

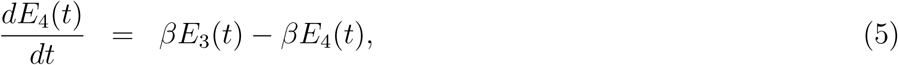

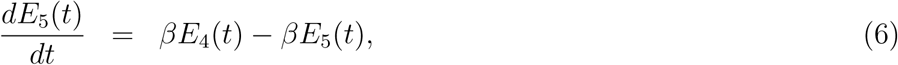

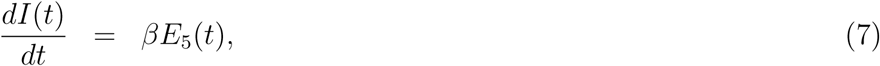

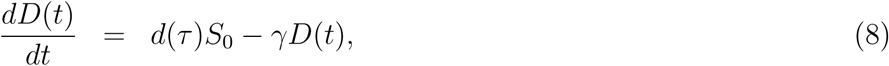

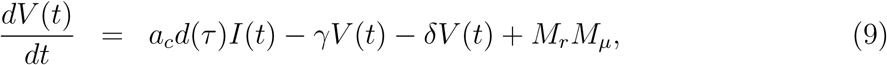

where

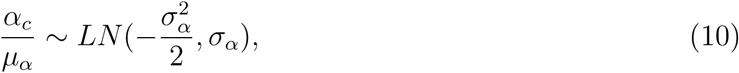

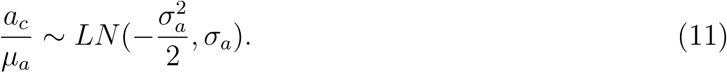

**Figure 1:**
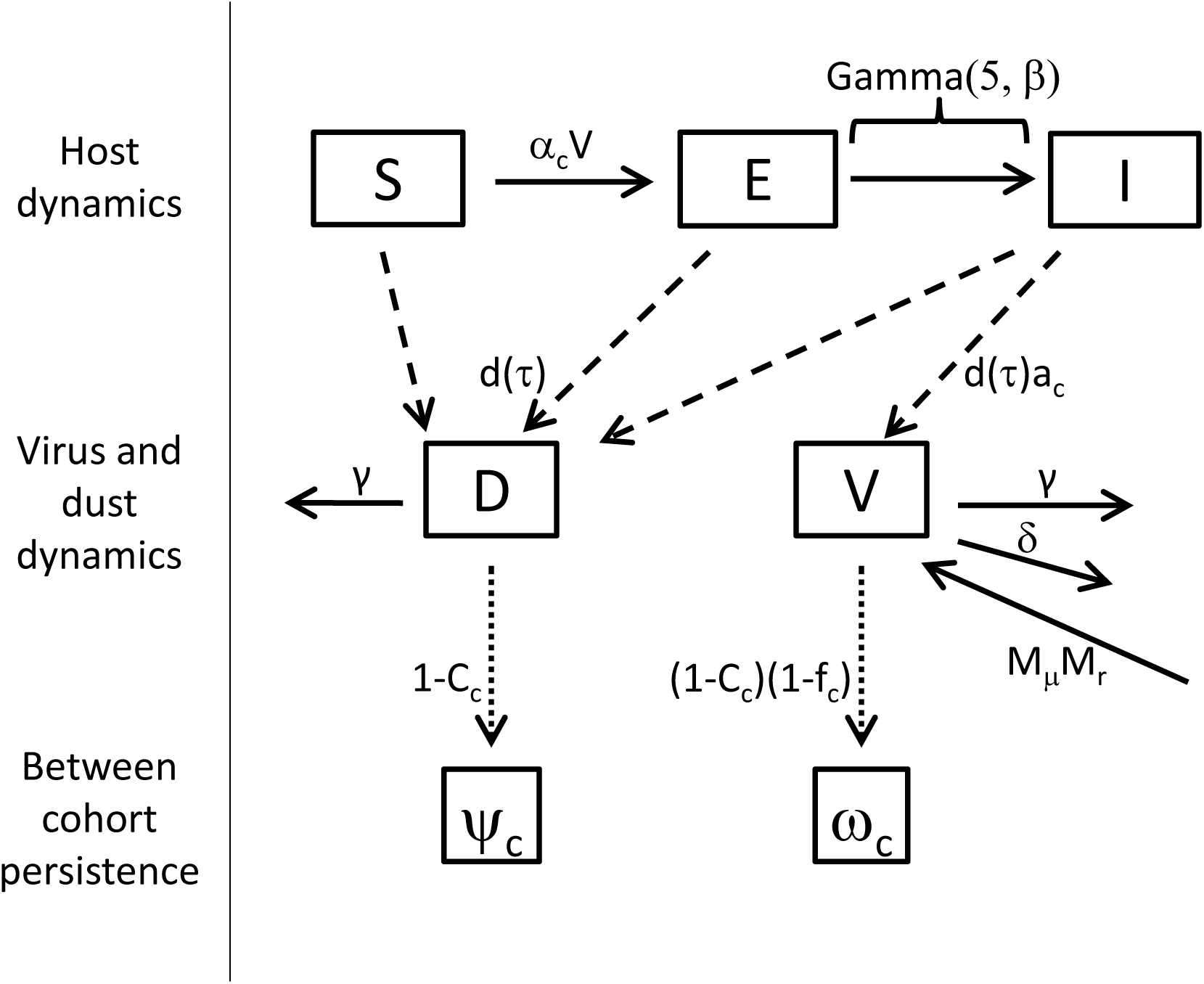
Schematic of the model. All states and parameters are as described in the main text. Solid lines indicate transitions between model classes. Dashed lines indicate that producing dust and virus does not cause birds to leave their current model class. Dotted lines indicate the between flock persistence of dust and virus. Note that without altering the model, we depict the exposed class as a single group, where the time until an exposed host becomes infected is gamma distributed with shape equal to 5 and rate equal to *β*.

In the above equations, *S*(*t*) is the number of susceptible birds at time *t. E*_*n*_(*t*) is the number of exposed birds in class *n* at time *t* where there are 5 total exposed classes (see below), *I*(*t*) is the number of infected birds at time *t, D*(*t*) is the total milligrams of dust in the house at time *t*, and *V*(*t*) is the total number of virus copies in the house at time *t. α*_*c*_ is the log normally distributed transmission rate of the virus in flock *c* determined by mean *μ*_*α*_ and adjusted scale parameter *σ*_*α*_, *β* is the transition rate between exposed classes, *γ* at which virus decays in a house. All birds in the house produce dust at a rate of *d*(*τ*), which is a function of flock age *τ*. Keeping in mind that bird mortality is assumed negligible, the total number of birds in the house at any time is equivalent to the initial number placed *SQ.* Infected birds from flock c also produce virus at a concentration of *a*_*c*_ per mg of dust produced, which is log normally distributed with mean *μ*_*a*_ and adjusted scale parameter *σ*_*a*_. Additionally, to allow for the possible reintroduction of virus on farms, a quantity of *M*_*μ*_ infectious virus copies are introduced at rate *M*_*r*_.

We model the exposed class as a series of compartments to create an incubation period of the virus that follows a gamma distribution (Wearing et al. 2005). In practice, we used *N* = 5 incubation classes with a transition rate *β* chosen to produce virus shed rates over time that were consistent with average viral copy number in feather tips of experimentally infected birds (Read et al. 2015), which is strongly correlated with viral shedding intensity (Baigent et al. 2013).

The above model describes disease dynamics within a flock of chickens, but a complete model of MDV dynamics must also incorporate the persistence of dust and virus between flocks. We follow the practice of nesting within flock disease dynamics into a discrete-time model to capture the multiple timescales of dynamics (Dwyer et al. 2000; Elderd et al. 2008). MDV is highly stable in the environment, and ventilation fans are typically off between flocks of chickens. As a consequence, the duration of downtime between flocks of birds is unlikely to strongly influence virus dynamics. Rather, the reduction in virus and dust between flocks is likely the result of cleaning. This cleaning process can be highly variable between flocks and between different farms. On some farms, bedding material, or “litter”, is reused for several flocks or up to several years. Between flocks on these farms, only caked or wet areas of litter are removed and top dressed with new litter. On other farms, litter is completely changed between each flock. Full removal of litter involves removing all bedding material down to the clay floor or cement floor. Also between flocks, farmers may or may not use forced air blowers to blow down chicken dust from fans, walls, and other above ground surfaces. Blow downs are often followed by a “thermal fog”, comprised of a vaporized disinfectant and an insecticide. Typically once per year, farmers will do a “wet clean and disinfect”, which is a wash down by spraying all surfaces with high pressure, low volume water. These wet cleans are typically followed by spraying with disinfectant.

The outcome of the cleanout process is that dust and virus might be removed concurrently through mechanical processes, or virus may degrade without the removal of dust due to the action of disinfectants. We thus model the initial virus and dust concentration in flocks using the following model:

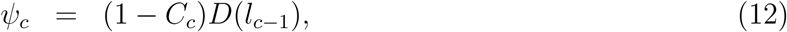

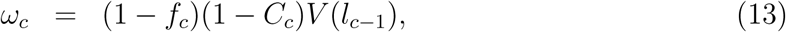
where

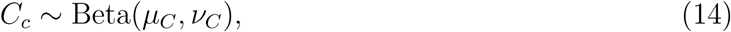

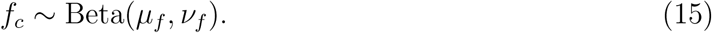

Here, *ψ*_*c*_ and *ω*_*c*_ are the respective total dust and total virus at the start time of flock *c. l*_*c*-1_ is the load out time of flock *c* − 1, making *D*(*l*_*c*-1_) and *V*(*l*_*c*-1_) the respective dust and virus at the end of the previous flock. *C*_*c*_ is the fraction of dust and virus removed between flocks *c*— 1 and c by the physical removal of dust, and *f*_*c*_ is the fraction of persisting virus that is removed between flocks *c* − 1 and *c* due to chemical disinfectants and other environmental decay factors. Note that *C*_*c*_ and *f*_*c*_ are beta distributed random numbers parameterized by their means (*μ*_*C*_, *μ*_*f*_) and sample sizes (*v*_*C*_, *v*_*f*_). The initial number of susceptible birds at the start of flock *c* is equal to the total number of birds placed in the house, and the initial number of exposed and infected birds are equal to 0. A list of all model parameters is provided in table 1.

**Table 1:** Parameter names, descriptions, and values

| Symbol | Description | Value (simulation) | Fixed | Source |
| --- | --- | --- | --- | --- |
| $\mu_\alpha$ | Transmission rate, mean | $1.652 \times 10^{-13}$ per susceptible bird per virus copy per day | Yes | (Atkins et al. 2011, 2013b) |
| $\sigma_\alpha$ | Transmission rate, scale | 0.3 | No | |
| $\mu_a$ | Virus shed rate, mean | $3 \times 10^6$ virus copies per infectious bird per day | No | |
| $\sigma_a$ | Virus shed rate, scale | 0.3 | No | |
| $\beta$ | Transition rate of exposed | 0.4 per exposed bird per day | Yes | (Read et al. 2015) |
| $d(\tau)$ | Dust shed rate at age $\tau$ | $10.8 + 368e^{-326/\tau^{1.64}}$ per bird per day | Yes | (Atkins et al. 2013a) |
| $\gamma$ | Ventilation rate | 0.2 per day | No | |
| $\delta$ | Within flock virus decay rate | 0.1 per day | No | |
| $\mu_C$ | Between flock cleanout, mean | 0.8 per virus copy or per mg of dust | No | |
| $\nu_C$ | Between flock cleanout, sample size | 1.0 | No | |
| $\mu_f$ | Between flock virus decay, mean | 0.8 per virus copy | No | |
| $\nu_f$ | Between flock virus decay, sample size | 1.0 | No | |
| $M_r$ | Virus reintroduction, rate parameter | 0.03 per day | No | |
| $M_\mu$ | Virus reintroduction, virus copies | $1 \times 10^8$ virus copies | No | |
| $S_0$ | Susceptible birds placed | 27000 birds | Yes | (Kennedy et al. 2017) |
| $V_0$ | Initial virus population size | $3 \times 10^{10}$ virus copies | No | |
| $D_0$ | Initial dust present | $1 \times 10^7$ mg | Yes | None |

To account for the fact that bird population sizes are finite, and for computational convenience, the above differential equations were replaced with their corresponding probabilistic transition equations (Kennedy et al. 2014, 2015a), where time was discretized to units of one day. We discretized all birds in the model to integer values, with transitions between classes determined by binomial probabilities. Virus and dust continued to be treated as continuous variables.

The above model allows for variation in virus transmission rate, virus shed rate from infected birds, dust and virus cleanout efficiency during downtime, virus decay rate during downtime, and virus reintroduction. These mechanisms for generating variation can however be removed from the model. Variation in virus transmission rate or in virus shed rate can be removed by respectively assuming that *σ*_*α*_ = 0 or *σ*_*a*_ = 0. Variation in the cleanout efficiency of dust and virus or in the decay rate of virus during downtime can be respectively removed by assuming that *v*_*C*_ or *v*_*f*_ are extremely large. Virus reintroduction can be removed by assuming that *M*_*r*_ = 0. Note that this additionally removes the parameter *M*_*μ*_ from the model. To test for the importance of these sources of variation in explaining virus dynamics, we therefore generated every version of the model that includes or excludes each of these factors (32 models in total), and we fit each of these models to data.

### Parameter estimation, model evaluation, and model comparison

To visualize the basic dynamics imposed by the model, we generated simulations of the most complex model using an illustrative parameter set (table 1), and we examined these simulations to identify common features in the model trajectories. We then used model inference to estimate parameters and determine how well the above model formulations describe data of MDV dynamics on commercial poultry farms. Several of our model parameters are already known from previous experiments. These include the rate of dust production as a function of bird age *d*(*τ*) (Atkins et al. 2013a), the incubation period of the virus (Islam and Walkden-Brown 2007; Read et al. 2015), which is determined by *β*, and the mean transmission rate of the virus *μ*_*α*_ (Atkins et al. 2011, 2013b). Placement dates, load out dates, and flock sizes were fixed at known values from the respective datasets being modeled. Point estimates of the remaining parameters were determined from the data using maximum likelihood.

The data used in this study are qPCR data reflecting the virus copy number (VCN) per mg of dust. These data were originally collected as part of a surveillance study quantifying the spatial and temporal variation of MDV across farms in Pennsylvania (Kennedy et al. 2017). We use two representative datasets from that study: the data from Farm A House 1, and the data from Farm E House 4 (fig. 2). These datasets were chosen because they were among the most exhaustively sampled farms and houses, and because they include highly disparate patterns of MDV dynamics. The Farm A House 1 dataset consists of 476 samples, collected at 163 time points. These data span 11 flocks with an average rearing duration of 84.3 days and an average of 27373 birds placed in each flock. Virus quantities on Farm A, although dynamic, remained at detectable levels throughout the study. The Farm E House 4 dataset consists of 454 samples, collected at 158 time points. These data span 21 flocks with an average rearing duration of 46.8 days and an average of 26855 birds placed in each flock. Virus quantities on Farm E went through periods of high virus concentration and periods where virus was undetectable.

**Figure 2:**
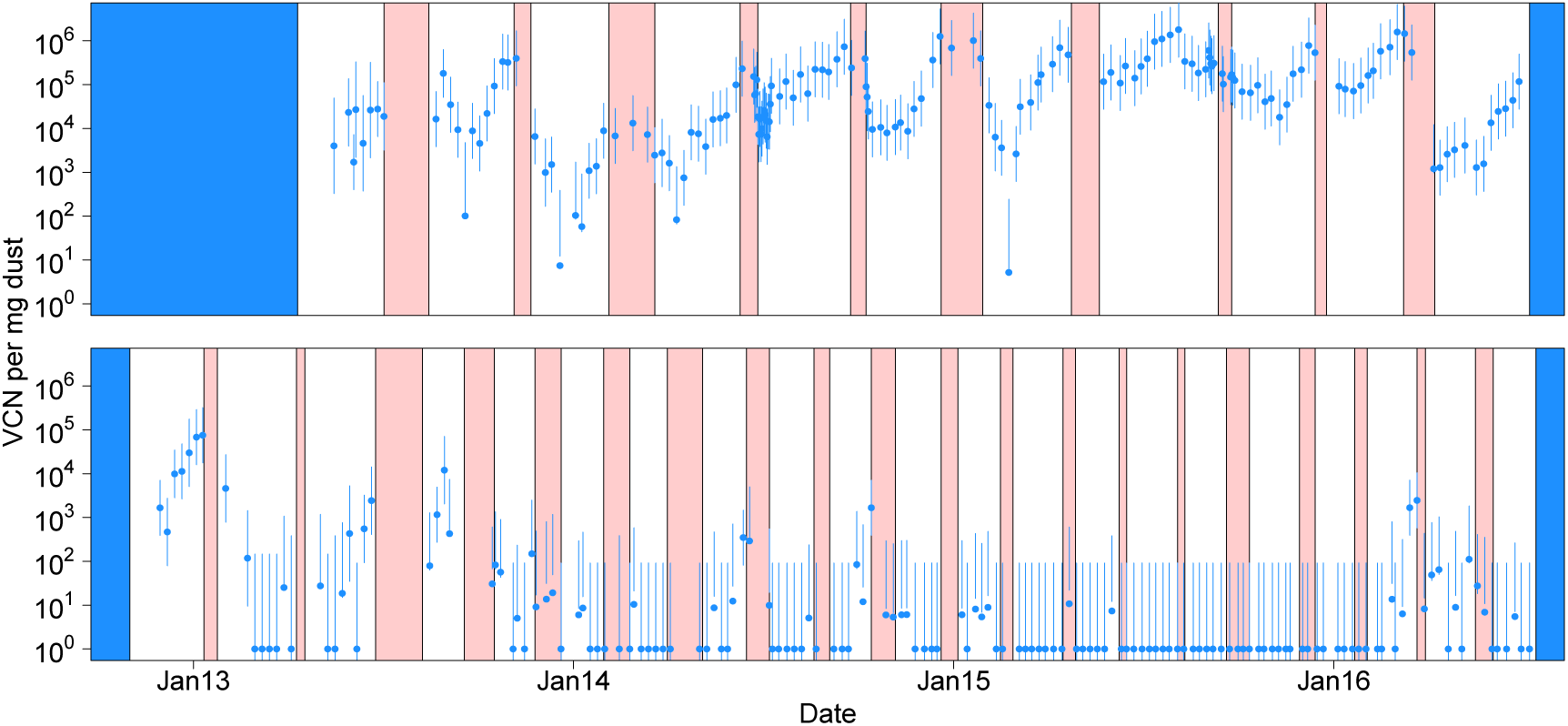
Data collected in Kennedy et al. (2017). The top panel shows the data for Farm A House 1, and the bottom panel shows the data for Farm E House 4. Red intervals show periods of down time, when birds are absent from houses. White intervals show periods when birds were present. Blue intervals show periods when surveillance data were not available. Points are the log mean plus one virus copy number per mg of dust collected at each sample time. Under the assumption that noise in the data is homoscedastic and normally distributed, bars show 95% confidence intervals around maximum likelihood estimates of virus copy number per mg of dust (Kennedy et al. 2017). Note that in rare circumstances, the error bars do not overlap the data points because the log mean virus concentration differs slightly from the maximum likelihood virus concentration. This discrepancy is partially due to the assumption of homoscedasticity in the data, an assumption that we relax to fit our models.

To construct a likelihood function, we first determined the marginal likelihood of virus being detectable by qPCR as a function of mean virus concentration. We treated 100 VCN per mg of dust as the limit of detection by qPCR, because below this concentration, virus detection is unreliable (Kennedy et al. 2017). Using the longitudinal data from Kennedy et al. (2017), we found that the probability of detection in biological replicates can be well described by a probit regression of the mean log_10_ VCN/mg of dust plus 1. Using a generalized linear model with binomial data and a probit link we found an intercept of −2.206 and a slope of 1.555 (fig. 3). For data that did not exceed our limit of detection, the likelihood was one minus the value resulting from this regression assuming the mean given by the model. For data that did exceed the limit of detection, the likelihood was the probability of detection multiplied by the probability of observing the particular virus concentration seen in the data (data were log_10_ plus 1 transformed), where the standard deviation was determined by a linear regression of the standard deviation with respect to the mean. We thus modeled this component of likelihood as a normal distribution with mean μ given by the model and standard deviation 1.127 – 0.151*μ* (fig. 3).

**Figure 3:**
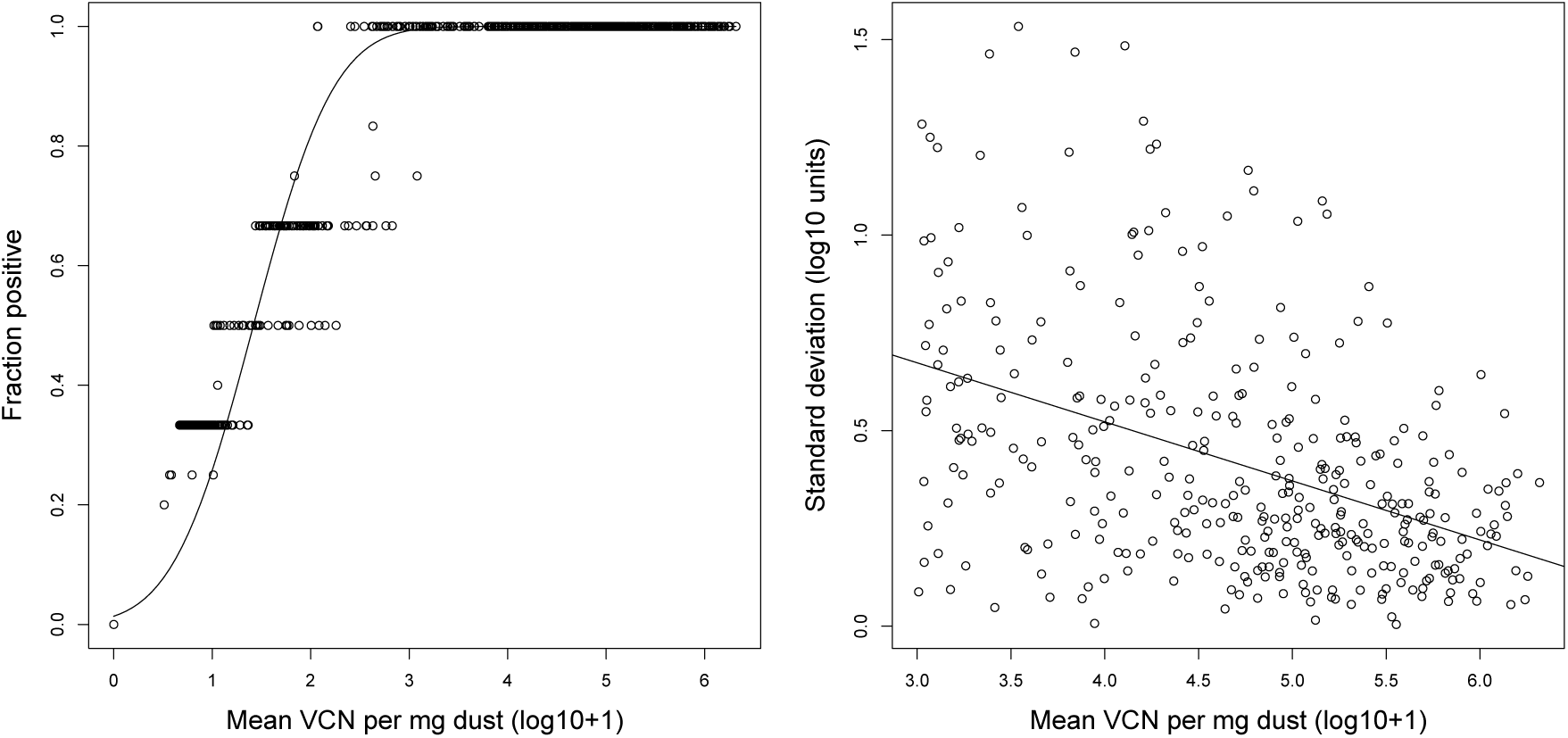
The empirically derived likelihood function using data from Kennedy et al. (2017). Shown on the left is the probability of virus detection across biological replicates as a function of measured log mean virus concentration. Points are the raw data, and the line is the best fit probit regression line. Shown on the right is the observed standard deviation of biological replicates from the same study, discarding when log mean virus concentrations were below 10^3^. Points are again the data, and the solid line is the best fit linear regression line.

Parameter estimation was performed using maximum likelihood methods with the ‘mif2’ function of the ‘pomp’ package in the R statistical computing language (King et al. 2016, 2017; R Core Team 2017). Fitting arguments used in ‘mif2’ included ‘Np=1000’, ‘Nmif=1500’, ‘cooling.fraction.50=0.5’, and ‘rw.sd=0.02’ for all regular parameters or ‘rw.sd=0.2’ for initial value parameters. Arguments ‘Np’, ‘Nmif’, and ‘cooling.fraction.50’ were modified for any model that repeatedly had trouble finding high likelihood parameter space. This was performed until at least 20 parameter sets were generated that had log likelihoods within 50 points of the current observed maximum likelihood. The likelihood of each final parameter set was determined using 10 iterations of ‘pfilter’ with ‘Np=10000’. All code and data are publicly available (github.com/dkenned1/KennedyDunnRead).

Models were compared using Akaike’s Information Criterion (AIC) (Burnham and Anderson 2002). To aid in comparison, we present ΔAIC and model weights (Burnham and Anderson 2002). We further explored the ability of the models to explain the data by simulating 5000 realizations from the weighted set of reasonable models, defined as any model with weight greater than 0.01. Each model was simulated using its respective maximum likelihood parameter estimates. We present envelopes that encompass central quantiles from these simulations.

MD takes four to twelve weeks post exposure to develop, and so the risk of leukosis condemnation is likely to positively correlate with the fraction of birds exposed to virus at least 30 days prior to load out. Hereafter, we refer to this fraction of birds exposed to virus 30 or more days before loadout as the “condemnation risk”. We present envelopes that encompass the fraction of birds exposed to MDV over time for the maximum likelihood estimated parameters from the set of reasonable models. We explore the impact of altering features of poultry rearing by exploring how two fold increases or decreases in model parameters relative to maximum likelihood estimates alter condemnation risk (rearing duration is altered by plus or minus 5 days).

To explore whether the parameter estimates that best explain the data from Farm A House 1 also provide a reasonable estimate for the data from Farm E House 4, and to explore the impact of rearing duration on MDV dynamics, we repeat these methods, applying the fitted models from Farm A House 1 to the rearing parameters (i.e. placement dates and load out dates) from Farm E House 4. We also do the reverse to see whether the best fit models and parameters from Farm E House 4 provide a reasonable explanation for the data from Farm A House 1.

## Results

The structure of the above model comes from a qualitative understanding of MDV natural history. Relating this qualitative understanding to quantitative dynamics, however, at a minimum requires simulation of the model with parameter values within their respective plausible ranges. In fig. 4, we show simulated trajectories of the model for conventional duration (40 days) and long duration (80 days) rearing periods using the illustrative parameter set from table 1. In fig. 5 we show the envelopes produced by simulating the model 5000 times. These two figures demonstrate at least five features of MDV dynamics that emerge from our mechanistic model. First, the sampled virus dynamics are inherently stochastic. This stochasticity can be visualized as the jaggedness in the curves, and it becomes less pronounced as virus concentrations increase (fig. 4). Second, different realizations of the model generate variable virus concentrations even when all simulations are performed with a single set of parameters (fig. 4). Third, virus densities within flocks tend to form “U” or “J” shaped trajectories, with virus concentrations starting relatively high, decreasing early on during the rearing period and increasing back to high concentrations later (fig. 5). Fourth, the sharpness of the “U” shaped trajectories vary between different flocks and realizations, sometimes generating less pronounced troughs than other times (fig. 4). Fifth, the rearing duration can strongly alter virus dynamics (fig. 5).

**Figure 4:**
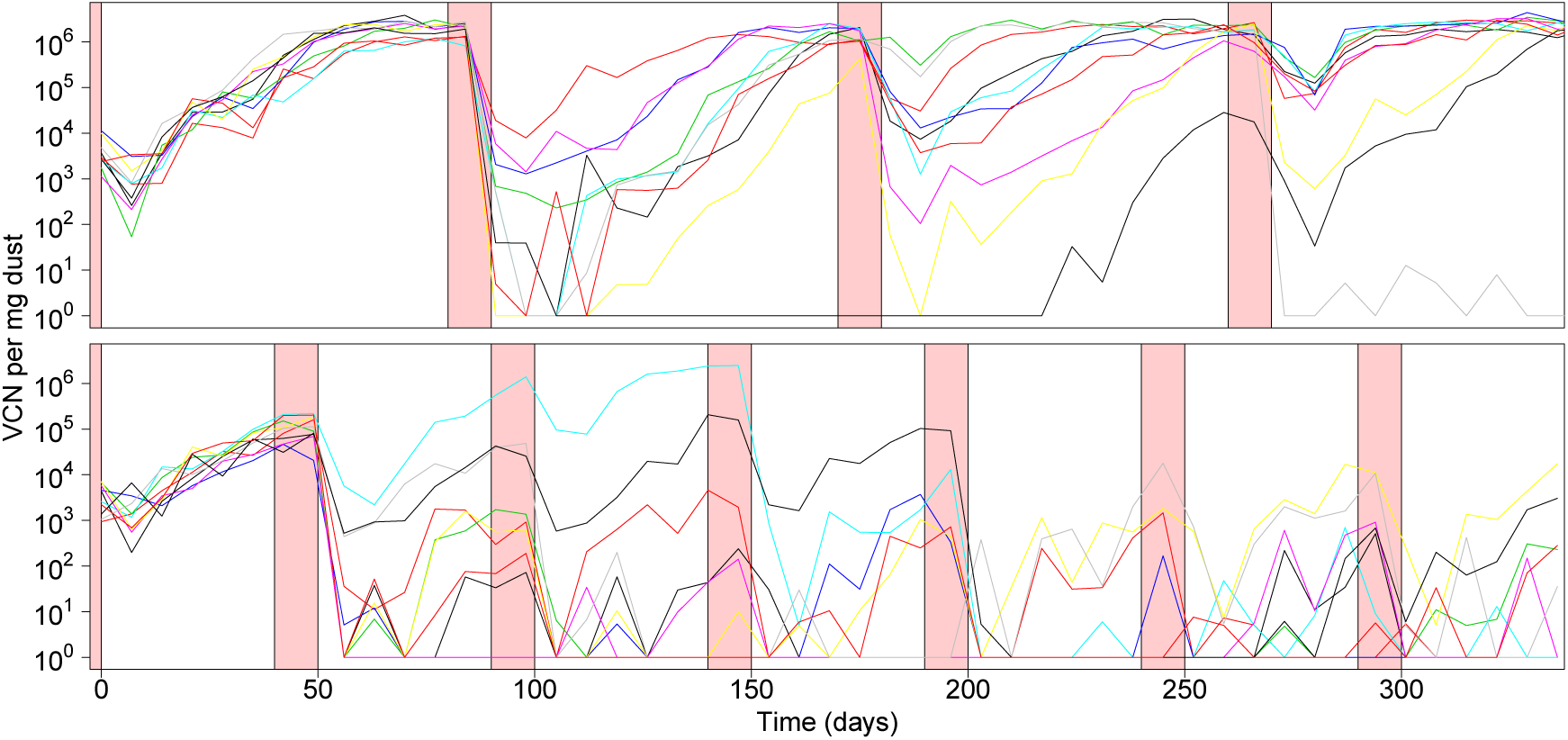
Ten simulated trajectories of the model shown in fig. 1 using an illustrative parameter set (table 1). As in fig. 2, red intervals show periods of down time, when birds are absent from houses. Both panels show trajectories using the same model parameters, but where rearing duration is 80 days (top) or 40 days (bottom). Note the differences in dynamics between these panels, as well as the differences in dynamics across trajectories and within trajectories between flocks.

**Figure 5:**
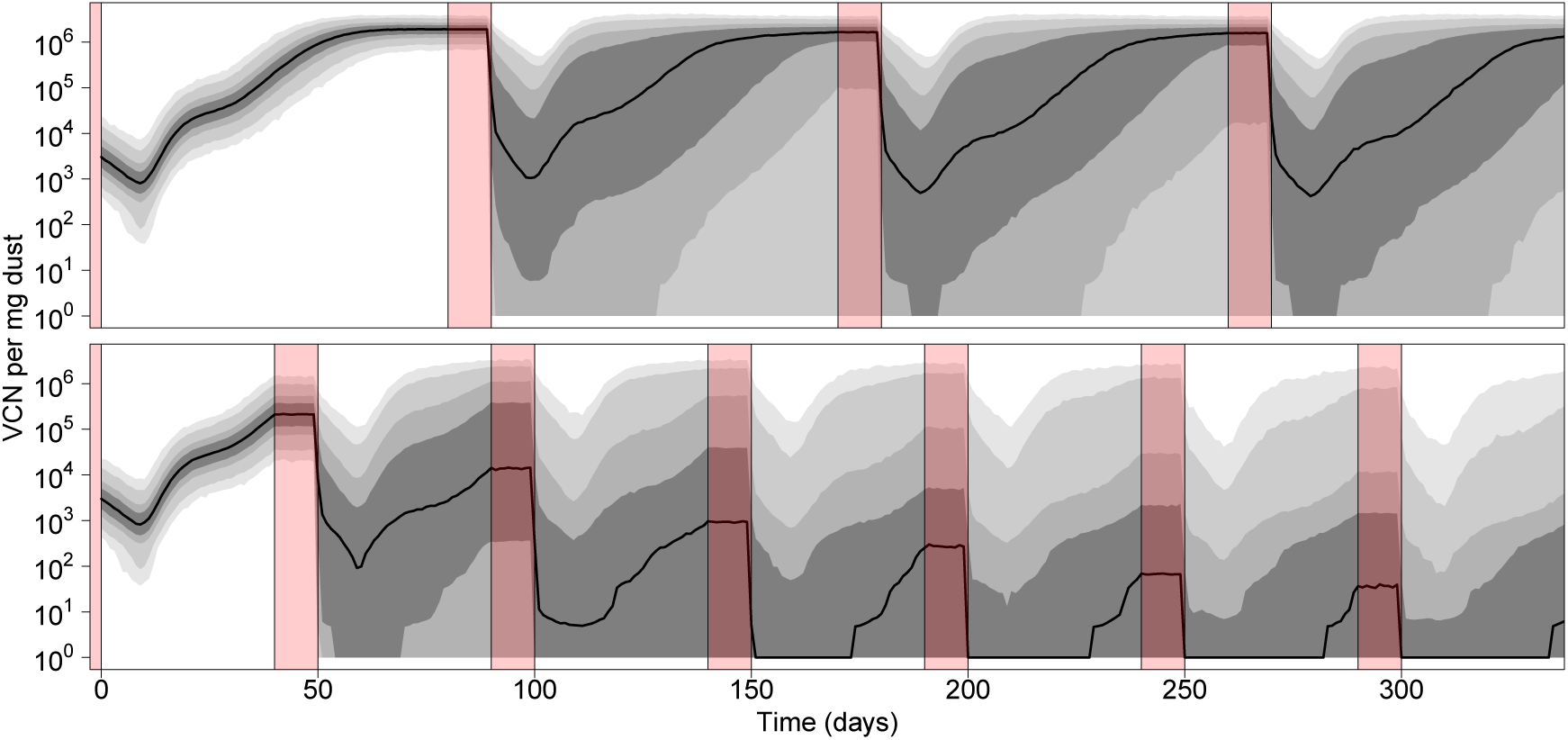
Envelope of model trajectories generated from 5000 simulations with the illustrative parameter set (table 1). Figure layout is identical that of fig. 4. Black solid lines show the median model prediction at each time point. Grey shaded regions show the respective 50%, 75%, 95% and 99% central quantiles.

By fitting our suite of models to the data from Farm A House 1, we found that three models have ΔAIC < 2, and are thus essentially indistinguishable (table 2). Each of these three best models include variation between flocks in the transmission rate of the virus *σ*_*a*_, in the shedding intensity of virus from infected birds *σ*_*α*_, and in the cleanout efficiency of dust and virus between bird flocks *v*_*C*_, suggesting that these features of the model are important to explaining virus dynamics. The decay of virus between flocks *v*_*f*_ and the reintroduction rate of virus *M*_*r*_ on the other hand are each excluded from at least one of the best three models, and therefore might not need to be included in models of virus dynamics on this farm.

**Table 2:** AIC table for Farm A House 1 dataset.

| Model | Variability parameters | MaxLHood | Params | ΔAIC | Weight |
| --- | --- | --- | --- | --- | --- |
| M31 | $\sigma_\alpha + \sigma_a + \nu_C + \nu_f + M_r$ | -500.1 | 12 | 1.5 | 0.22 |
| M30 | $\sigma_\alpha + \sigma_a + \nu_C + \nu_f$ | -511.2 | 10 | 19.7 | 0.00 |
| M29 | $\sigma_\alpha + \sigma_a + \nu_C + M_r$ | -500.3 | 11 | 0.0 | 0.47 |
| M28 | $\sigma_\alpha + \sigma_a + \nu_C$ | -502.7 | 9 | 0.8 | 0.31 |
| M27 | $\sigma_\alpha + \sigma_a + \nu_f + M_r$ | -507.2 | 11 | 13.8 | 0.00 |
| M26 | $\sigma_\alpha + \sigma_a + \nu_f$ | -520.7 | 9 | 36.7 | 0.00 |
| M25 | $\sigma_\alpha + \sigma_a + M_r$ | -535.4 | 10 | 68.1 | 0.00 |
| M24 | $\sigma_\alpha + \sigma_a$ | -615.8 | 8 | 225.0 | 0.00 |
| M23 | $\sigma_\alpha + \nu_C + \nu_f + M_r$ | -509.4 | 11 | 18.1 | 0.00 |
| M22 | $\sigma_\alpha + \nu_C + \nu_f$ | -515.4 | 9 | 26.1 | 0.00 |
| M21 | $\sigma_\alpha + \nu_C + M_r$ | -511.8 | 10 | 21.0 | 0.00 |
| M20 | $\sigma_\alpha + \nu_C$ | -510.3 | 8 | 13.9 | 0.00 |
| M19 | $\sigma_\alpha + \nu_f + M_r$ | -536.2 | 10 | 69.7 | 0.00 |
| M18 | $\sigma_\alpha + \nu_f$ | -524.0 | 8 | 41.3 | 0.00 |
| M17 | $\sigma_\alpha + M_r$ | -551.1 | 9 | 97.6 | 0.00 |
| M16 | $\sigma_\alpha$ | -611.8 | 7 | 214.9 | 0.00 |
| M15 | $\sigma_a + \nu_C + \nu_f + M_r$ | -535.4 | 11 | 70.2 | 0.00 |
| M14 | $\sigma_a + \nu_C + \nu_f$ | -534.9 | 9 | 65.1 | 0.00 |
| M13 | $\sigma_a + \nu_C + M_r$ | -514.8 | 10 | 27.0 | 0.00 |
| M12 | $\sigma_a + \nu_C$ | -531.8 | 8 | 56.9 | 0.00 |
| M11 | $\sigma_a + \nu_f + M_r$ | -538.0 | 10 | 73.4 | 0.00 |
| M10 | $\sigma_a + \nu_f$ | -558.1 | 8 | 109.5 | 0.00 |
| M9 | $\sigma_a + M_r$ | -570.0 | 9 | 135.4 | 0.00 |
| M8 | $\sigma_a$ | -699.6 | 7 | 390.6 | 0.00 |
| M7 | $\nu_C + \nu_f + M_r$ | -580.0 | 10 | 157.4 | 0.00 |
| M6 | $\nu_C + \nu_f$ | -576.0 | 8 | 145.4 | 0.00 |
| M5 | $\nu_C + M_r$ | -592.6 | 9 | 180.6 | 0.00 |
| M4 | $\nu_C$ | -598.1 | 7 | 187.6 | 0.00 |
| M3 | $\nu_f + M_r$ | -604.4 | 9 | 204.1 | 0.00 |
| M2 | $\nu_f$ | -615.3 | 7 | 221.9 | 0.00 |
| M1 | $M_r$ | -684.6 | 8 | 362.6 | 0.00 |
| M0 | None | -997.1 | 6 | 983.6 | 0.00 |

We find somewhat different results when we examine the Farm E House 4 data. While we again find that several models provide comparable fits to the data according to AIC (table 3), we find that for this dataset, every model that includes reintroduction of virus (*M*_*μ*_ and *M*_*r*_) has a better AIC score than every model that excludes this mechanism. This result suggests that virus reintroduction is important to explaining dynamics on this farm. In addition, we find that virus reintroduction alone is unable to explain the breadth of variation present on the farm, as noted by the fact that model M1 which includes only stochastic virus introduction is not in our set of models with weights greater than 0.01. Rather, we find that each of the models that provide reasonable explanations for the data also include variation between flocks in either virus transmission rate or virus shed rate from infected birds.

**Table 3:** AIC table for Farm E House 4 dataset.

| Model | Variability parameters | MaxLHood | Params | ΔAIC | Weight |
| --- | --- | --- | --- | --- | --- |
| M31 | $\sigma_\alpha + \sigma_a + \nu_C + \nu_f + M_r$ | -361.5 | 12 | 3.9 | 0.03 |
| M30 | $\sigma_\alpha + \sigma_a + \nu_C + \nu_f$ | -375.9 | 10 | 28.7 | 0.00 |
| M29 | $\sigma_\alpha + \sigma_a + \nu_C + M_r$ | -361.9 | 11 | 2.7 | 0.05 |
| M28 | $\sigma_\alpha + \sigma_a + \nu_C$ | -376.9 | 9 | 28.7 | 0.00 |
| M27 | $\sigma_\alpha + \sigma_a + \nu_f + M_r$ | -362.3 | 11 | 3.3 | 0.04 |
| M26 | $\sigma_\alpha + \sigma_a + \nu_f$ | -377.1 | 9 | 29.0 | 0.00 |
| M25 | $\sigma_\alpha + \sigma_a + M_r$ | -362.5 | 10 | 1.8 | 0.08 |
| M24 | $\sigma_\alpha + \sigma_a$ | -385.4 | 8 | 43.7 | 0.00 |
| M23 | $\sigma_\alpha + \nu_C + \nu_f + M_r$ | -362.4 | 11 | 3.7 | 0.03 |
| M22 | $\sigma_\alpha + \nu_C + \nu_f$ | -378.1 | 9 | 30.9 | 0.00 |
| M21 | $\sigma_\alpha + \nu_C + M_r$ | -362.4 | 10 | 1.6 | 0.08 |
| M20 | $\sigma_\alpha + \nu_C$ | -377.7 | 8 | 28.2 | 0.00 |
| M19 | $\sigma_\alpha + \nu_f + M_r$ | -362.4 | 10 | 1.6 | 0.08 |
| M18 | $\sigma_\alpha + \nu_f$ | -381.3 | 8 | 35.4 | 0.00 |
| M17 | $\sigma_\alpha + M_r$ | -363.7 | 9 | 2.3 | 0.06 |
| M16 | $\sigma_\alpha$ | -388.7 | 7 | 48.2 | 0.00 |
| M15 | $\sigma_a + \nu_C + \nu_f + M_r$ | -361.9 | 11 | 2.7 | 0.05 |
| M14 | $\sigma_a + \nu_C + \nu_f$ | -376.1 | 9 | 27.0 | 0.00 |
| M13 | $\sigma_a + \nu_C + M_r$ | -361.6 | 10 | 0.0 | 0.19 |
| M12 | $\sigma_a + \nu_C$ | -376.2 | 8 | 25.2 | 0.00 |
| M11 | $\sigma_a + \nu_f + M_r$ | -361.7 | 10 | 0.3 | 0.16 |
| M10 | $\sigma_a + \nu_f$ | -377.8 | 8 | 28.5 | 0.00 |
| M9 | $\sigma_a + M_r$ | -362.7 | 9 | 0.3 | 0.16 |
| M8 | $\sigma_a$ | -386.7 | 7 | 44.1 | 0.00 |
| M7 | $\nu_C + \nu_f + M_r$ | -369.5 | 10 | 15.9 | 0.00 |
| M6 | $\nu_C + \nu_f$ | -415.2 | 8 | 103.2 | 0.00 |
| M5 | $\nu_C + M_r$ | -368.6 | 9 | 12.1 | 0.00 |
| M4 | $\nu_C$ | -417.3 | 7 | 105.4 | 0.00 |
| M3 | $\nu_f + M_r$ | -369.8 | 9 | 14.5 | 0.00 |
| M2 | $\nu_f$ | -453.1 | 7 | 177.0 | 0.00 |
| M1 | $M_r$ | -370.4 | 8 | 13.7 | 0.00 |
| M0 | None | -524.3 | 6 | 317.5 | 0.00 |

In fig. 6 we show the parameter estimates that emerged as the maximum likelihood estimates from the set of reasonable models as defined by having model weights greater than 0.01. Using these parameter estimates in simulating our model we generate envelopes that shows the variation in the data expected given the model structure and maximum likelihood parameter sets. These simulations incorporate both process error (variation in virus dynamics between simulated trajectories) and observation error (variation in the data due to measurement error). Note the differences in the envelopes that are generated from these different datasets (fig. 7). On Farm A House 1, virus concentrations generally remain at high levels across many different realizations. On Farm E House 4 on the other hand, the virus concentrations quickly fall to low levels, but with reasonable chances for outbreaks to reemerge in future flocks due to virus reintroduction. These patterns are similarly reflected in the data. In fig. 8, we show how the fraction of birds exposed to virus changes over time for the two farms using maximum likelihood estimates from the set of reasonable models, revealing that substantially more birds are exposed to virus on Farm A House 1 than on Farm E House 4.

**Figure 6:**
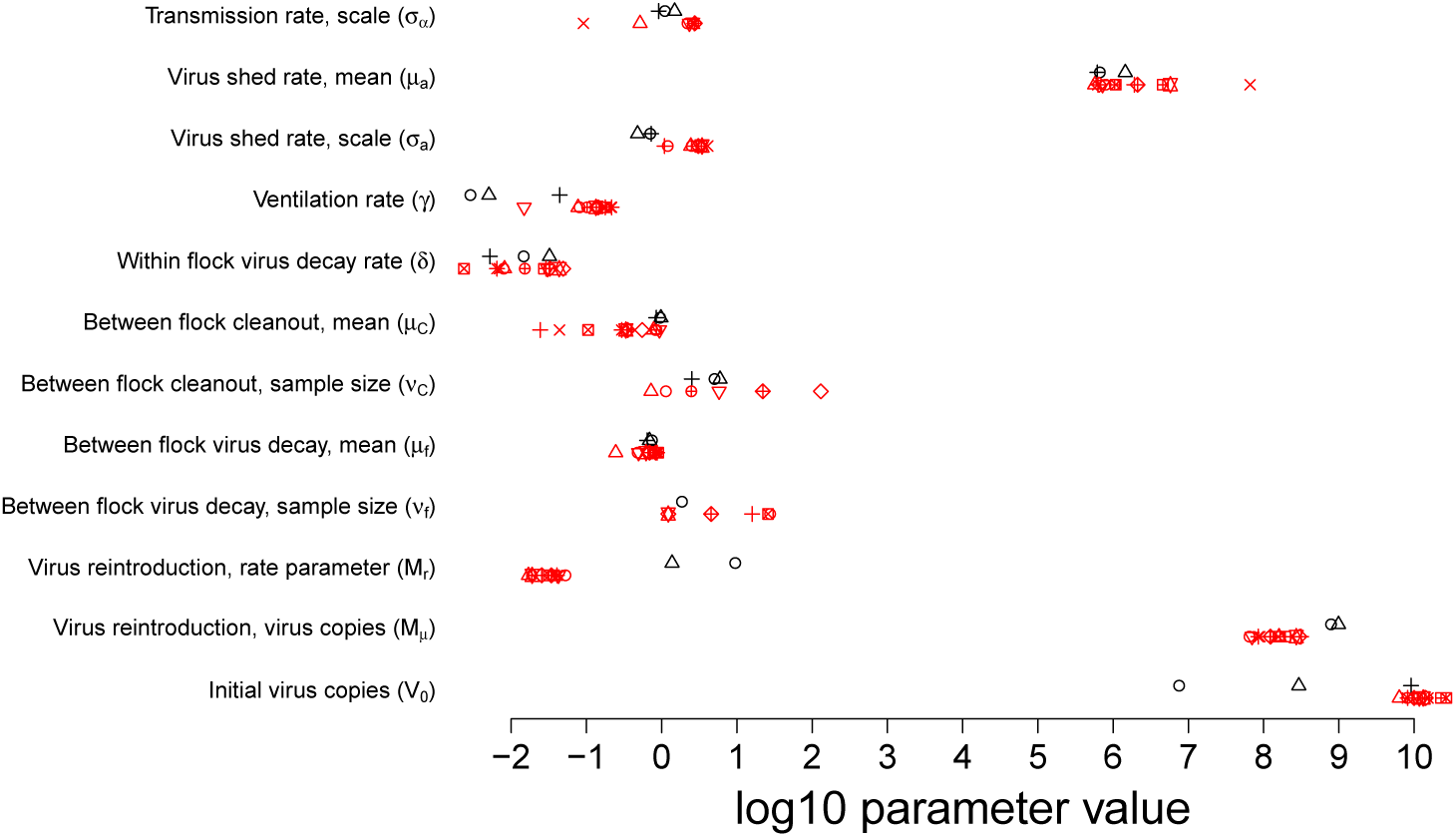
Maximum likelihood parameter estimates for each of the models with AIC weights greater than 0.01. Shown in black are the models fit to the Farm A House 1 data. Shown in red are the models fit to the Farm E House 4 data. Each shape shows a different model. For parameters that are missing from a particular model, no point is shown.

**Figure 7:**
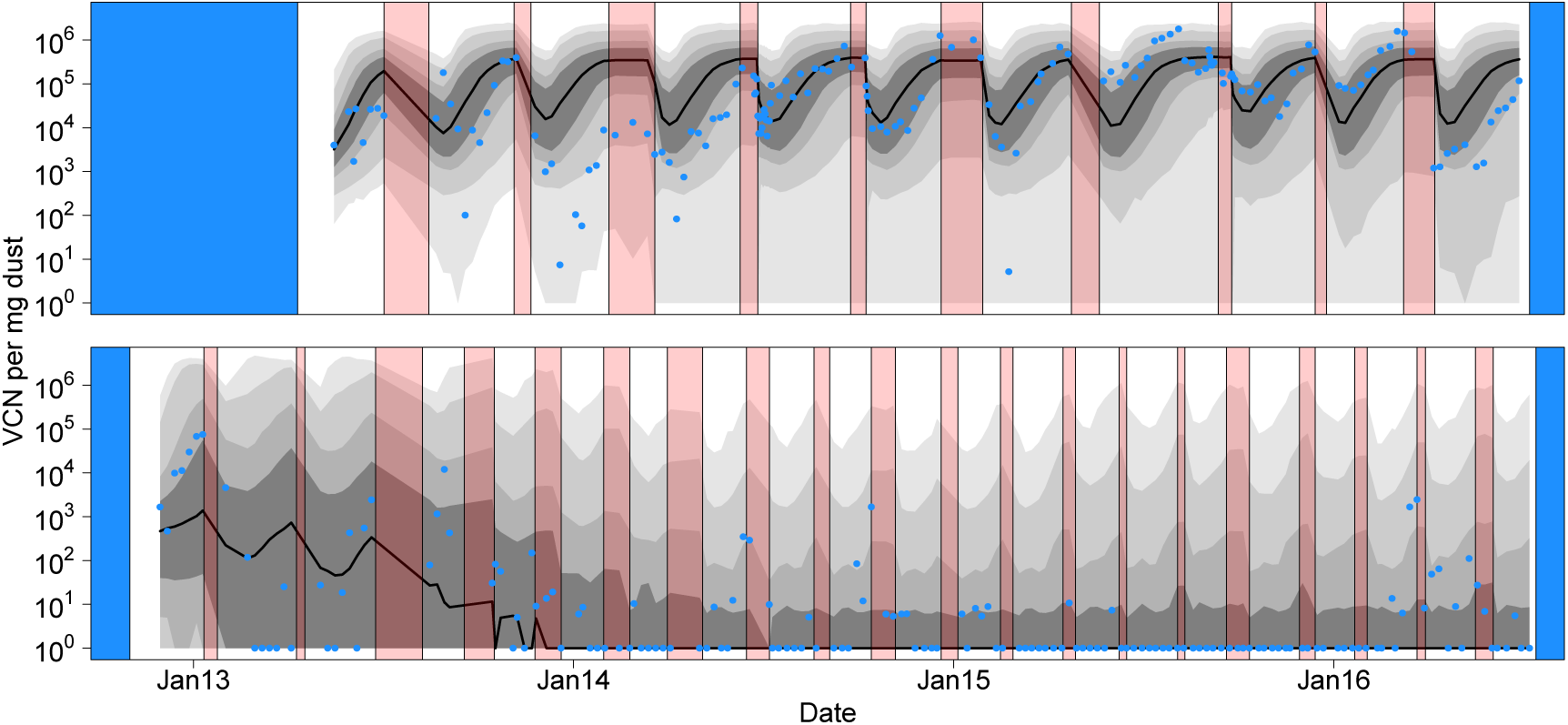
Fit of the model to the data. Red intervals as in fig. 4 denote periods of farm down time when birds were absent. Blue intervals denote periods where surveillance was not conducted. As in fig. 5, grey intervals of different darkness show 50%, 75%, 95%, and 99% central quantiles for the model predictions, with a solid black line depicting the median. Quantiles were generated from 5000 model realizations, where the model used to generate each realization was randomly selected based on AIC weights. Blue points denote the log mean virus concentration calculated from the data (Kennedy et al. 2017). The top panel shows the results for Farm A House 1 and the bottom panel shows the results from Farm E House 4. Tick marks on the x-axis denote the beginning of each new calendar year.

**Figure 8:**
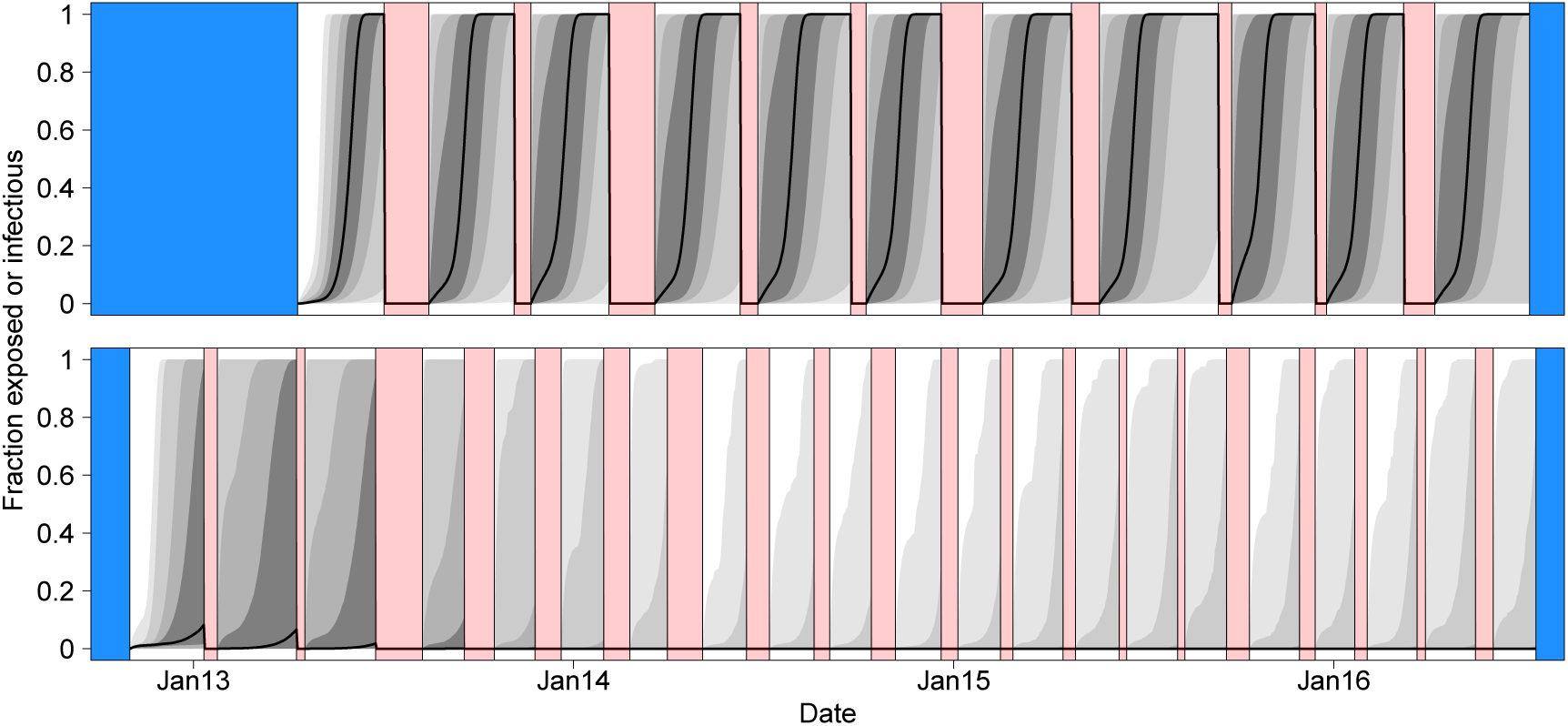
Fraction of birds present in a house that are currently in the exposed or infectious class. Note that we arbitrarily set the value to zero when birds were absent from the house. See fig. 7 for interpretation of colors and lines.

Fig. 9 shows how changes in farming practices (i.e., changes in parameter values of the model) are predicted to impact condemnation risk. Note that despite large differences between the two farms in overall condemnation risk, altering most parameters has similar effects on both farms. The transmission rate scale *σ*_*α*_, the mean virus shed rate *μ*_*a*_, the virus shed rate scale *σ*_*a*_, the mean between flock cleanout efficiency *μ*_*C*_ and the number of birds placed *S*_0_ have relatively large effects on condemnation risk for both farms. The ventilation rate *γ*, the sample size of cleanout efficiency *ν*_*C*_, and the mean decay rate of virus between flocks *μ*_*f*_ have relatively large impacts on only one of the farms. The decay rate of virus during the rearing period *δ*, the sample size of virus decay between flocks *v*_*f*_, the rate of virus reintroduction *M*_*r*_, the quantity of virus reintroduced *M*_*μ*_, and the initial virus population size *V*_0_ have relatively little effect on the condemnation risk for either farm.

**Figure 9:**
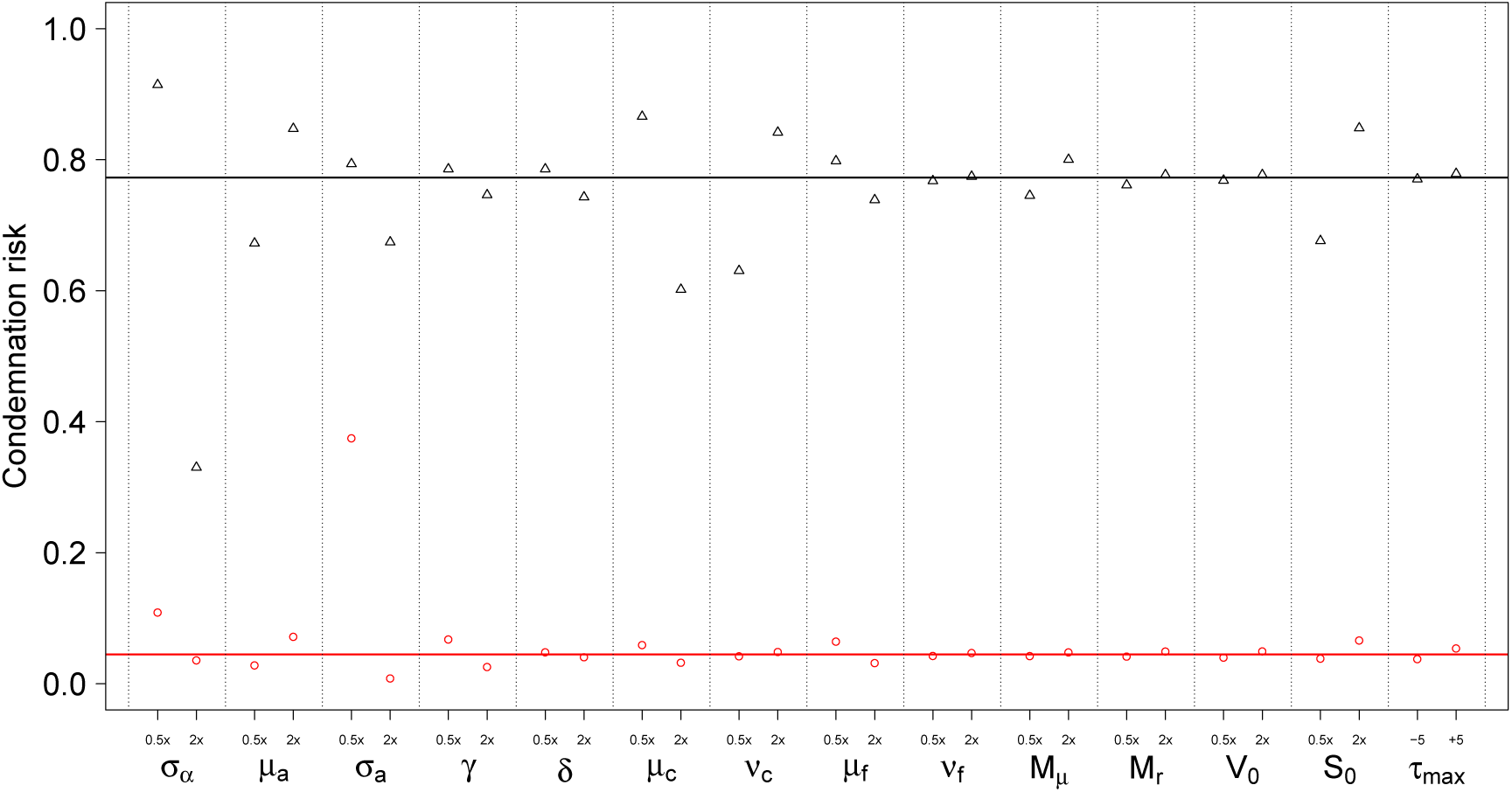
Simulated condemnation risk for Farm A House 1 (black) and Farm E House 4 (red) when altering the model parameters (solid horizontal lines mark condemnation risk at maximum likelihood estimates). “0.5x” on the x-axis denotes halving the parameter value, and “2x” denotes doubling the parameter value, with the exception of the mean cleanout efficiency *μ*_*C*_ and the mean virus degradation between flocks *μ*_*f*_ For these latter parameters “0.5x” is the case where twice as much virus or dust is held over between flocks, bounded to be non-negative (i.e. argmax(2/*μ* − 1, 0)), and “2x” is where hold over virus or dust is cut in half (i.e. 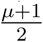). Respectively, −5 and +5 denote decreasing or increasing the rearing duration *τ*_max_ by 5 days. Note that the direction of the effect for each parameter is always the same between Farm A House 1 and Farm E House 4, but some parameters have disproportionately more effect on the condemnation risk of one farm than the other.

Applying our maximum likelihood parameter estimates from Farm A House 1 to the rearing conditions of Farm E House 4, and vice versa, allows us to visualize the relative impacts of differences in model parameters and differences in rearing duration on virus dynamics. Fig. 10 shows that the maximum likelihood parameter estimates inferred from virus concentration data on one of these farms do not accurately predict virus concentrations on the other farm, which can be seen in the mismatch between the prediction envelopes and the data. Comparisons between figs.7 and 10 show that if Farm A House 1 were to shorten rearing durations to those of Farm E House 4, it would lead to an approximate 10 fold reduction in median virus concentration. If Farm E House 4 were to extend rearing durations to those of Farm A House 1, it would lead to an increase of median virus concentrations from undetectable levels to between 10^2^ and 10^3^ virus copies per mg of dust.

**Figure 10:**
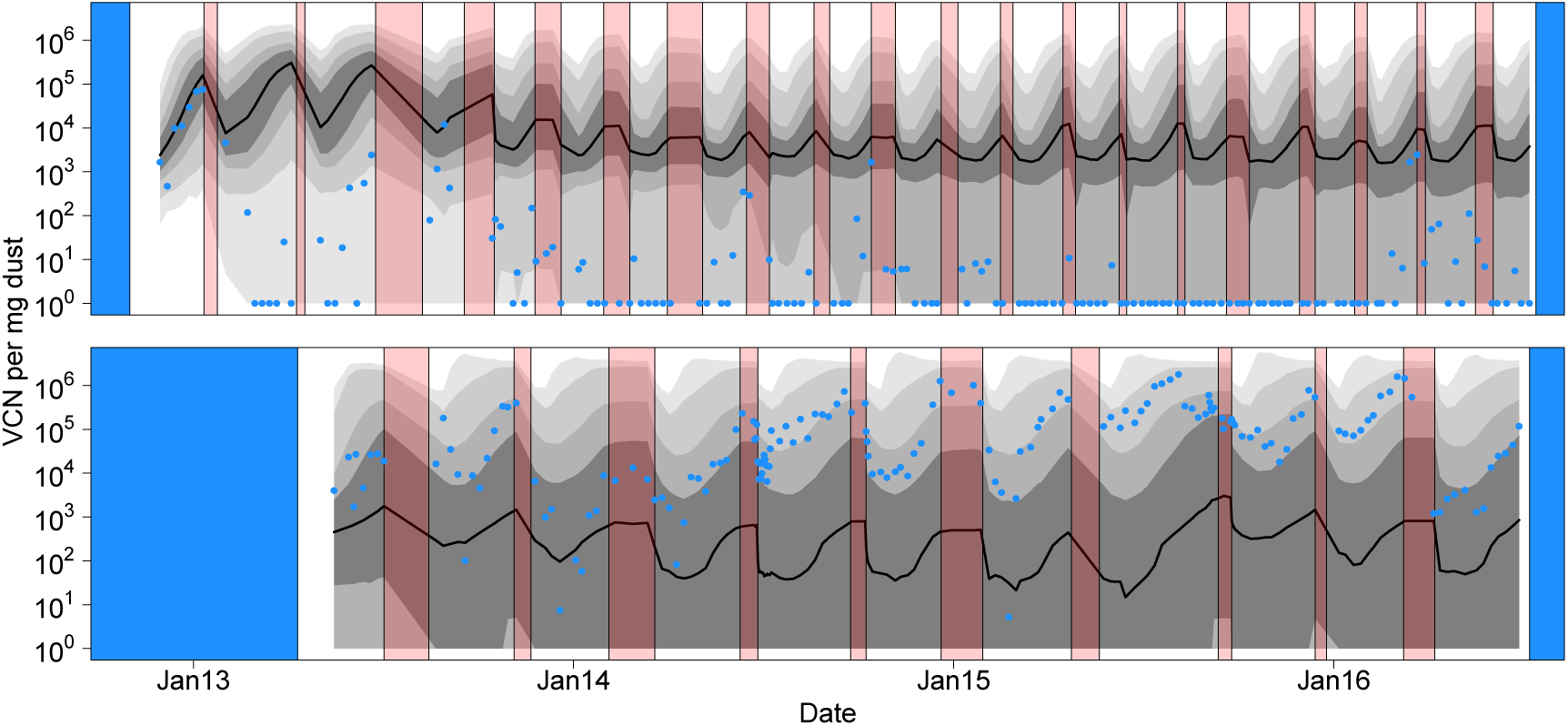
Fit of the model to the data when model weights and parameter estimates from Farm A House 1 are applied to Farm E House 4 (top), and the reverse (bottom). Note that the panels are swapped relative to fig. 7 to facilitate comparison of the impact of rearing duration on virus dynamics. The lack of agreement between model predictions and data suggest that the parameter values governing virus transmission differ at least slightly between these farms.

## Discussion

Here, we constructed mechanistic models to describe MDV dynamics across multiple flocks of chickens reared on a commercial chicken farm. We used data on virus concentration to identify the mechanisms that drive variation in virus dynamics between farms (tables 2 and 3), and to provide the first data-inferred estimates of the parameters that define these mechanisms (fig. 6). Our models were able to capture the dynamics of two different datasets, one in which virus intensities tended to be much higher than in the other (figs. 2 and 7). The different dynamics in these two datasets arise from slight differences in model structure, parameter estimates, and rearing practices. We then showed that altering the parameters of these models influences condemnation risk (fig. 9), giving insight into how MDV losses might be managed should virus evolution undermine existing vaccines.

Two previous sets of models have been developed to describe MDV dynamics. The first set by Atkins and colleagues (Atkins et al. 2013a,b) uses an individual-based approach that captures many of the biological details particular to MDV dynamics. The second set by Rozins and Day (Rozins and Day 2016, 2017) uses an impulsive-differential-equation based approach that allows the models to be completely communicated through a set of equations. Our modeling approach is a combination of these two approaches. Like the Rozins and Day approach, our models have an underlying impulsive differential equation structure, making it easy to describe them. Like the models of Atkins and colleagues, our models allow for stochasticity and biological details that are not captured by simpler models. The key difference between the approach presented here and previous approaches, however, is that the parameters of our model have been determined by statistical integration with field data. Our best models are therefore those which have withstood confrontation with data, and these models are therefore more likely to produce accurate predictions than untested models.

Previous surveillance on MDV dynamics has revealed that virus concentrations in dust tend to form U-shaped trajectories (Kennedy et al. 2017). This pattern was attributed to dilution of virus early in the rearing period when few birds are shedding virus-contaminated dust followed by increasing concentration of virus later in the rearing period when many birds are shedding virus-contaminated dust (Kennedy et al. 2017). Here, we have shown that this qualitative shape naturally arises from the structure of our models (fig. 5), and that our models quantitatively explain such patterns in field data (fig. 7).

We used model comparison methods to identify the features of poultry farming that lead to variation in MDV dynamics between flocks and between farms. For the Farm A House 1 data, we found that all reasonable models included flock-to-flock variation in virus transmission rates *σ*_*α*_, virus shedding rates *σ*_*a*_, and clean out efficiency between flocks *v*_*C*_, thus highlighting the importance of these mechanisms for explaining dynamics on this farm (table 2). This result suggests that flocks of birds differ in their susceptibility to virus and in their shedding rate should they become infected. It also suggests that clean out efficiency varies on this farm and this variation has had detectable effects on virus dynamics. For the Farm E House 4 data, we found that all reasonable models included flock-to-flock variation in either virus transmission rates *σ*_*α*_ or virus shedding rates *σ*_*a*_, and they also all included the stochastic reintroduction of virus *M*_*μ*_ and *M*_*r*_ (table 3). This again suggests that flock-to-flock variation in either disease susceptibility or virus shedding is affecting virus dynamics. Our finding that virus is stochastically reintroduced on this farm, also suggests that eradication of virus may be quite challenging. Even if a control program were capable of clearing infection from a farm, there appears to be substantial risk of reintroduction, meaning that control measures might need to be sustained indefinitely to maintain an infection-free farm.

Although our two datasets identified different mechanisms as key to driving MDV dynamics, this should not be surprising. Such an outcome might arise from true differences in the mechanisms that generate variation between farms, but it also might arise due to other differences in the data. For example, the importance of stochastic reintroduction of virus is likely realized on Farm E House 4, because virus concentrations are low in that dataset. Similar levels of virus reintroduction might also occur when virus is common such as on Farm A House 1, but because virus is common, this mechanism would have only a negligible impact on the dynamics of the virus. This outcome highlights that the absence of a mechanism in a reasonable model does not mean that the mechanism is not acting, but only that it is not important to explaining those particular data. Of the mechanisms tested by our approach, variable decay of virus between flocks *v*_*f*_ is the only mechanism that was not necessary to explain either of the datasets studied.

Examination of the maximum likelihood parameter estimates that arise from the sets of reasonable models reveals that there is substantial overlap of the range of these maximum likelihood parameter estimates across the datasets (fig. 6). There are only three parameters where the ranges of the maximum likelihood estimates do not clearly overlap between the two farms. These are the virus shed rate scale *σ*_*a*_, the virus reintroduction rate parameter *M*_*r*_ and the virus reintroduction quantity parameter *M*_*μ*_. We note that variation in virus shed rate *σ*_*a*_ might be hard to distinguish from virus reintroduction on Farm E House 4, leading to an overestimate of this parameter. Likewise, the rate and quantity of virus reintroduction might be hard to quantify on Farm A House 1 because these parameters are likely obscured by the high virus concentrations on this farm. Both lines of reasoning are supported by the observations that 4 out of 12 of the reasonable models for Farm E House 4 do not include the parameter *σ*_*a*_, and 1 out of the 3 reasonable models for Farm A House 1 do not include the parameters *M*_*r*_ and *M*_*μ*_, implying a great deal of uncertainty regarding the point estimates of these parameters.

The parameter estimates that arose from model fitting give further insight into model reliability, virus management, and perhaps even virus evolution. Regarding model reliability we can compare our parameter estimates to those measured in previous studies. To our knowledge, the only parameter in our model that was directly measured through lab experiments is the mean virus shed rate *μ*_*a*_. Two studies have provided estimates of this value from bivalent vaccinated chickens. Islam and Walkden-Brown (2007) found that birds shed 6.36 log_10_ virus copies per mg of dust. Atkins et al. (2011) tested three virus strains of difference virulence rank and found respective shedding rates of 6.04, 7.06, and 7.12 log_10_ virus copies per mg of dust. Our estimates (mean=6.30, s.d.=0.61 log_10_ virus copies per mg of dust) is in close agree with these previous studies. The decay rate of virus *δ* was never quantitatively measured, but two studies have looked at the infectiousness of dust after storage at room temperature. Jurajda and Klimes (1970) found no change in virus infectivity after 44 days and Witter et al. (1968) found that 8 of 14 infectious dust samples maintained infectivity after 112 days. Our median estimate of *δ* implies a half life of viral persistence of 66 days (range 20 to 425 days), which appears to be on similar scale to that of these previous experiments, further supporting that our model is reasonably capturing the biology of the system.

Our estimates of virus reintroduction rate *M*_*r*_ on Farm E House 4 imply a median reintroduction probability of 3.2% per day (range 1.7% to 5.1%). This equates to approximately one virus reintroduction event per month. Based on these estimates, a typical flock of chickens is likely to be exposed to virus even if the virus is not present at the time of placement. However, the timing of virus reintroduction is an important factor in determining whether virus amplification will take place. Introductions late in the rearing cycle, for example, are unlikely to amplify to high levels before birds are removed for processing, highlighting the importance of good biosecurity early in the rearing cycle. Nevertheless, the frequent reintroduction rate suggests that maintaining the absence of virus is likely to require rearing conditions that limit the long term amplification of virus.

Our estimates of the parameters dictating between flock removal of virus and dust *μ*_*C*_ and decay of virus *μ*_*f*_ suggest that the cleanout and sanitization processes between flocks are fairly efficient, with the median estimate for cleanout efficiency at 95%, and the median estimate of virus decay at 69% (note that the median for *μ*_*C*_ includes only the estimates from Farm A House 1 because the estimates of this parameter from Farm E House 4 are highly uncertain across models). This relatively high efficiency implies that current practices are good at reducing dust and virus between flocks. This efficiency, however, may be a cause for concern given that Rozins and Day (2017) found efficient cleaning between flocks can promote the evolution of increased MDV virulence by decreasing the lifespan of the pathogen and thus decreasing the cost of virulence.

Nevertheless, we found that condemnation risk is highly sensitive to the mean cleanout efficiency *μ*_*C*_ and flock size *S*_0_ when compared against the impact of the mean virus decay between flocks *μ*_*f*_, the ventilation rate *δ*, or the within flock virus decay rate *δ*. This suggests that, should vaccine efficacy be reduced by pathogen evolution, condemnation rates might best be managed by focusing efforts on blowing down dust, changing litter between flocks, and reducing flock sizes, as opposed to improving chemical treatments between flocks or increasing ventilation rates within flocks. Reducing the frequency *M*_*r*_ or intensity *M*_*μ*_ of virus reintroduction from other farms also appears to have little influence on condemnation risk. Nevertheless, *M*_*r*_ would become a key parameter should the goal change from reducing to eliminating MD related condemnation.

We also found that condemnation risk is highly sensitive to the transmission rate scale parameter *σ*_*α*_, and the virus shed rate scale parameter *σ*_*a*_. Because the scale parameters *σ*_*α*_ and *σ*_*a*_ are directly related to variability between flocks in transmission rate and virus shed rate, our analysis demonstrates that condemnation risk is highest when flocks have little variability between them in transmission rate and virus shed rate. This observation highlights the value of intervening when a farm is having problems, even if the intervention is economically unsustainable in the long term, because a reduction in the transmission and virus shedding rates within a single flock can have large impacts on overall condemnation risk. Currently, such a strategy is practiced through methods such as reactive Rispens vaccination in response to MD breaks. Cycling flocks of chickens between standard broilers and MD-resistant broiler might have similar benefits (Hunt and Dunn 2013).

Our results highlight the importance of rearing duration on virus dynamics. Long rearing durations give the virus ample time to spread through a flock, and reach high concentrations in dust (figs. 4 and 10). The historical decline in MDV-associated condemnation is strongly correlated with declines in rearing durations, making it difficult to tease apart the relative impact that rearing duration and other changes in biosecurity and vaccine administration have had on disease control (Kennedy et al. 2015b). Nevertheless, our model shows that rearing duration is one of several factor that can potentially be manipulated to control problems with disease should they reappear.

Given the history of pathogen evolution undermining vaccine efficacy, any long term control strategy for MDV must carefully consider the ecology of disease transmission, and its impact on pathogen evolution. Our hope is that the data-tested models that we present here will lay the ground work for this approach. Future work might use these models to ask how altering rearing practices, approximated by changing model parameters, affects the profitability of poultry farming as well as how this might alter competition between virus strains and in turn the evolution of the virus.

## Acknowledgments

This work was funded by the Institute of General Medical Sciences (R01GM105244), National Institutes of Health and United Kingdom Biotechnology and Biological Sciences Research Council as part of the joint NSF-NIH-USDA Ecology and Evolution of Infectious Diseases program, and by the RAPIDD program of the Science and Technology Directorate, Department of Homeland Security and Fogarty International Center, National Institutes of Health (DAK, AFR). The funders had no role in study design, data collection and analysis, decision to publish, or preparation of the manuscript.

